# Assessing the feasibility of machine learning for ancient DNA age prediction: limitations and insights

**DOI:** 10.1101/2025.11.09.687346

**Authors:** M. Kazanskii, M. Golikova, A. Kasianov, A. Ivanova, A. Suraganov, L. Uroshlev

## Abstract

We investigated the possibility of estimating the age of ancient biological samples directly from their DNA damage profiles using supervised machine learning. Traditional dating methods such as radiocarbon dating, dendrochronology rely on either material context or isotope composition, while our approach exploits intrinsic molecular degradation signatures. Using damage statistics obtained from ancient DNA sequencing data, we trained several regression models to predict sample age over a temporal range of up to 10,000 years. Despite initial correlations between specific damage features and age, cross-validation and external testing revealed no statistically significant predictive signal beyond mean-based baselines. These findings indicate that, in the current formulation, DNA damage information alone is insufficient for reliable age estimation. However, this negative result provides important methodological insight: environmental and biochemical factors appear to dominate damage variation, effectively masking chronological signal. We suggest that integrating contextual data, expanding labeled datasets, and incorporating physical models of DNA decay may improve future attempts. Our study thus contributes to a transparent assessment of the limitations and prospects of DNA-based fossil dating.

## Introduction

Since its emergence, ancient DNA (aDNA) research has transformed archaeology and comparative anthropology. Modern sequencing techniques now make it possible to recover and analyze DNA from samples over one million years old, with the current record reaching approximately two million years for environmental DNA and about 1.3 million years for individual organisms. These advances allow researchers to identify the origins of ancient remains and compare them both with one another and with contemporary populations. aDNA studies have even led to the discovery of previously unknown human species, such as the Denisovans Slon et al. [2017].

Despite this progress, several challenges in aDNA data analysis remain unresolved. One of the most prominent is the ability to accurately estimate the age of a biological sample solely from its DNA sequence.

Under normal physiological conditions, random DNA damage is continuously corrected by cellular repair machinery. However, after death, these repair processes cease, initiating the first phase of DNA degradation. Intracellular nuclease are released as cellular structures break down. Once liberated, these enzymes gain access to the genome and begin cleaving long DNA molecules into shorter fragments. This process progressively reduces the genome into increasingly smaller DNA fragments Dabney et al. [2013].

Once the initial hydrolytic breakdown of DNA subsides, the molecule begins to accumulate mutations, including those driven by environmental factors such as ultraviolet radiation. Two prominent chemical processes occur during this stage: depurination and cytosine deamination.

In depurination, the N-glycosidic bond linking a purine base (adenine or guanine) to the sugar–phosphate backbone is cleaved, resulting in loss of the base and potentially causing a base skip during sequencing. In cytosine deamination, the amino group is removed from cytosine, converting it into uracil. Because standard high-throughput DNA sequencing does not faithfully read uracil residues, this modification appears in sequencing data as a cytosine-to-thymine substitution Briggs et al. [2010].

In addition to these modifications, DNA can undergo further damage related to exposure to reactive oxygen species or the external environment. The older the sample, the more damage it accumulates. Therefore, using features such as DNA molecule length, number of substitutions, and mutations, it is possible to construct a model for estimating the age of a sample. Although there is currently no single model of DNA degradation over time Allentoft et al. [2012]. Moreover, environmental factors greatly influence DNA preservation Poinar & Stankiewicz [1999]. But there is evidence of a correlation between features such as DNA fragmentation and sample age.

However, developing a model for age estimation presents several challenges. One major source of complexity is the influence of environmental conditions. Optimal preservation requires low temperatures, dry conditions, minimal exposure to ultraviolet radiation, and limited oxygen availability. In contrast, DNA degradation accelerates in environments characterized by high or fluctuating temperatures, elevated humidity or frequent precipitation, direct sunlight, acidic or highly alkaline soils, and intense microbial activity Bollongino et al. [2008].

It should also be noted that only a small number of studies have focused on investigating the dynamics of nuclear DNA decay. Much more often, authors focus on mitochondrial or bacterial DNA. The rate of nuclear DNA decay is twice that of mitochondrial DNA decay, which is why most authors prefer to work with mitochondria.

In this work, we investigate whether the chronological age of ancient biological samples can be inferred directly from their DNA damage patterns using supervised machine learning. Motivated by known associations between postmortem molecular decay processes and sample age, we extract features such as fragment length distributions and characteristic substitution patterns from ancient DNA sequencing data and evaluate multiple regression approaches. Our goal is to assess the feasibility and limitations of DNA-based temporal prediction and determine whether damage signatures contain a stable signal that can serve as a proxy for sample age independent of contextual or isotopic information.

## Methods

### Data

The age prediction for ancient DNA samples was formulated as a regression task. For model training, we used output tables generated by DamageProfiler Neukamm et al. [2021], which report the frequency of each mutation type. DamageProfiler is a tool designed to characterise nucleotide misincorporation patterns in ancient DNA. It provides detailed profiles of post-mortem DNA damage, identifying and quantifying substitution types that are characteristic of ancient samples. In particular, it detects deamination-driven changes such as C→T and G→A transitions, which typically occur near the fragment ends. The transition mutation types considered in this study are as follows:

- TOTAL: Total number of nucleotides at the specified positions.
- A, C, G, T: Counts of individual nucleotides observed at the specified positions.
- *G → A, C → T*, *A → G, T → C, A → C, A → T*, *C→ G, C→ A, T→ G, T→ A, G→ C, G → T*: Single-base substitutions representing transitions or transversions from one nucleotide to another. *A→ −*, *T→ −*, *C→ −*, *G→ −*: Deletions where a nucleotide is lost from the sequence.
- →*A*, →*T*, →*C*, →*G*: Insertions where a nucleotide is added to the sequence.
- →*S*: Soft-clipped bases—nucleotides located at the ends of reads that do not align to the reference genome.

As the dataset, we used the Allen Ancient Genome Diversity Project Reich et al. [2024] (hereafter referred to as the Harvard dataset), which contains 216 BAM files. This large-scale ancient DNA sequencing initiative aims to explore genetic diversity, identify population-specific SNPs, and reconstruct human migration patterns. The sequences in this dataset have a median coverage of approximately 5×, and the sample ages range from roughly 500 to nearly 12,000 years. The distribution of the ages of the samples is presented on Figure 1. All reads were generated on Illumina platforms, with a mean read length of 60 bp, followed by basecalling and alignment to the hs37d5 reference genome. The hs37d5 (hg19) reference genome was chosen because it remains widely used in ancient DNA studies, ensuring compatibility with legacy datasets and tools. GRCh38, while more recent, introduces alternate haplotypes and coordinate changes that complicate alignment consistency and batch-effect control. Most samples underwent UDG treatment (uracil-DNA glycosylase) to remove uracils and improve sequencing accuracy. Additionally, a standardized bioinformatic pipeline was applied across all samples, resulting in a highly uniform and consistent dataset that is well suited for our model training.

**Figure 1:**
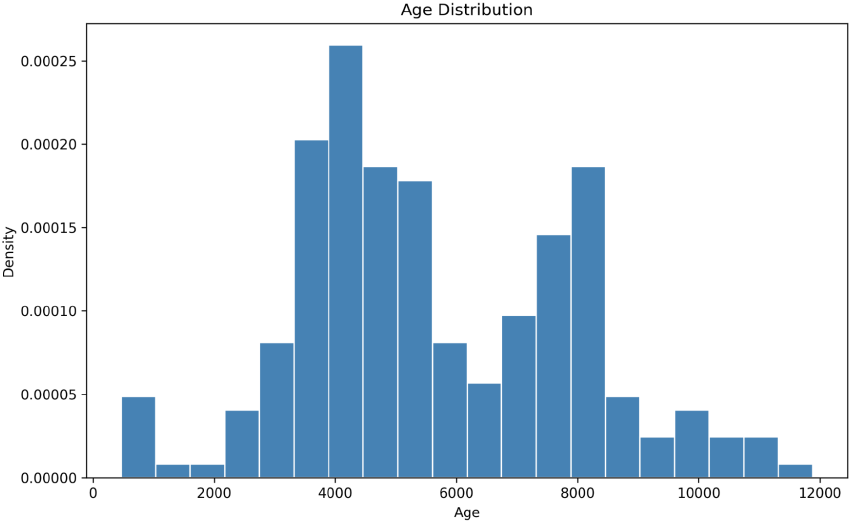
The distribution of samples in Harvard dataset by ages.

**Figure 2:**
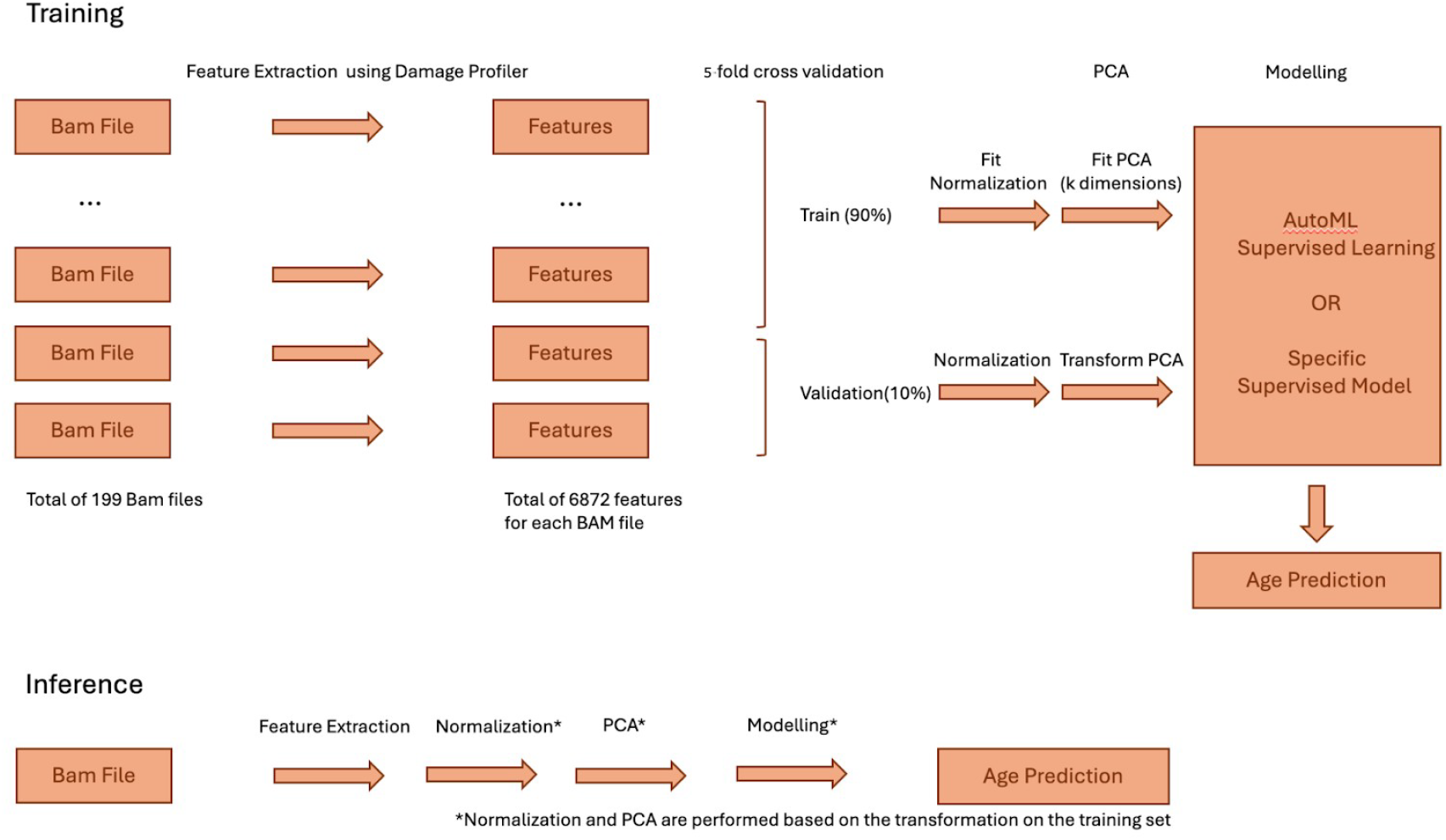
The general pipeline for the Age Prediction.

In addition to the Harvard dataset, we collected an independent set of publicly available ancient genomes from multiple published studies spanning Europe, Asia, Africa, and the Americas (listed below). These samples were downloaded from the European Nucleotide Archive (ENA) and the NCBI Sequence Read Archive (SRA), and they include individuals ranging from approximately 350 to approximately 10,850 years in age. We used this additional dataset as an external test. The detailed list of samples, including extracted ages, study names, and corresponding references, is provided in Appendix 1.

We performed clustering on the DamageProfiler output and visualized the data in a two-component space for the combined dataset (Harvard dataset samples and the external ctest set). The Harvard samples formed two distinct clusters, while the external test samples formed several additional clusters. This clear separation between sample groups suggests the presence of batch effects.

To mitigate batch effects Leek et al. [2010], we grouped all samples according to their publication of origin, resulting in a total of 15 groups. We then partitioned the dataset into train, validation, and test subsets such that each group was assigned exclusively to one subset. This strategy ensured that no publication group was shared across splits. Notably, when samples were not grouped in this manner, the model initially appeared to perform well; however, it was learning to predict the source publication rather than the sample age.

The data from Damage Profiler has been divided into [Train & Validation — Test] datasets with proportion [0.9 — 0.1]. The Train & Validation subset was used further in the pipeline with the cross-validation. The Mean Average Error (MAE) and R-squared have been used as a metric for the performance. In addition for the inference we used the BAM files not included in the Harvard dataset files so that we showed that the algorithm is generalized well beyond the training set (no significant batch effect present). Also we showed the current limitations of the algorithm for the inference data.

### Pipeline

We calculated the statistics for each type of mutation transition and the transition probability for each sequence length (the typical output of the DamageProfiler file Neukamm et al. [2021]).

A total of 199 BAM files were used in this study. The original Harvard dataset contained 216 BAM files, but 17 were excluded due to memory constraints associated with processing individual files. Each BAM file yielded approximately 5,900 features, corresponding to all combinations of mutation types and positional sequence contexts. Columns containing NaN values (400 columns) were removed. Additionally, DamageProfiler features corresponding to read positions greater than 60 (out of 100) were excluded. This step is particularly important for inference, as low read coverage in ancient DNA increases noise at later read positions. Notably, removing these positions did not substantially affect model performance.

Feature normalization was performed as follows: counts of individual nucleotides (e.g., “A”, “G”) and the “S” feature were normalized by the “TOTAL” read count; mutation transition features (e.g., “A→T”, “A→?”) were normalized by the count of the originating nucleotide (“A” in this example); and reverse transitions (e.g., “→A”) were normalized by the count of the target nucleotide. Normalization was applied row-wise to the DamageProfiler output. Finally, the “TOTAL” column was removed to prevent the model from exploiting coverage-related correlations rather than true temporal signal.

The problem occurs as one of *p << n* type of problems Hastie & Tibshirani [2003], or where the number of features is more than the number of data points. Therefore, some kind of dimensionality reduction is necessary in order to predict the age of the sample. We used the PCA algorithm Jolliffe & Cadima [2016] for this purpose. The usage of PCA is justified in our case since the features are highly correlated. The output number of features was a hyper-parameter for a pipeline (from 10 to 120). Before feeding the data to the PCA, each feature has been normalized to [0, 1]. The general pipeline for age prediction is schematically presented in 2.

In this study, we evaluated multiple regression models to identify the most effective approach and corresponding hyperparameters for ancient sample age prediction. We tested a range of supervised learning algorithms (summarized in Figure 3). Models implemented via the Scikit-learn library Pedregosa et al. [2011] included Ridge Regression, Lasso, ElasticNet, Random Forest, Support Vector Regression (SVR), k-Nearest Neighbors (KNN), Bayesian Ridge, and Decision Trees. In addition, we incorporated the XG-Boost algorithm using the XGBoost framework Chen & Guestrin [2016]. Model performance was compared against a baseline predictor equal to the mean age of the training set. To ensure reliable evaluation, all models were assessed using 5-fold cross-validation Browne [2000].

**Figure 3:**
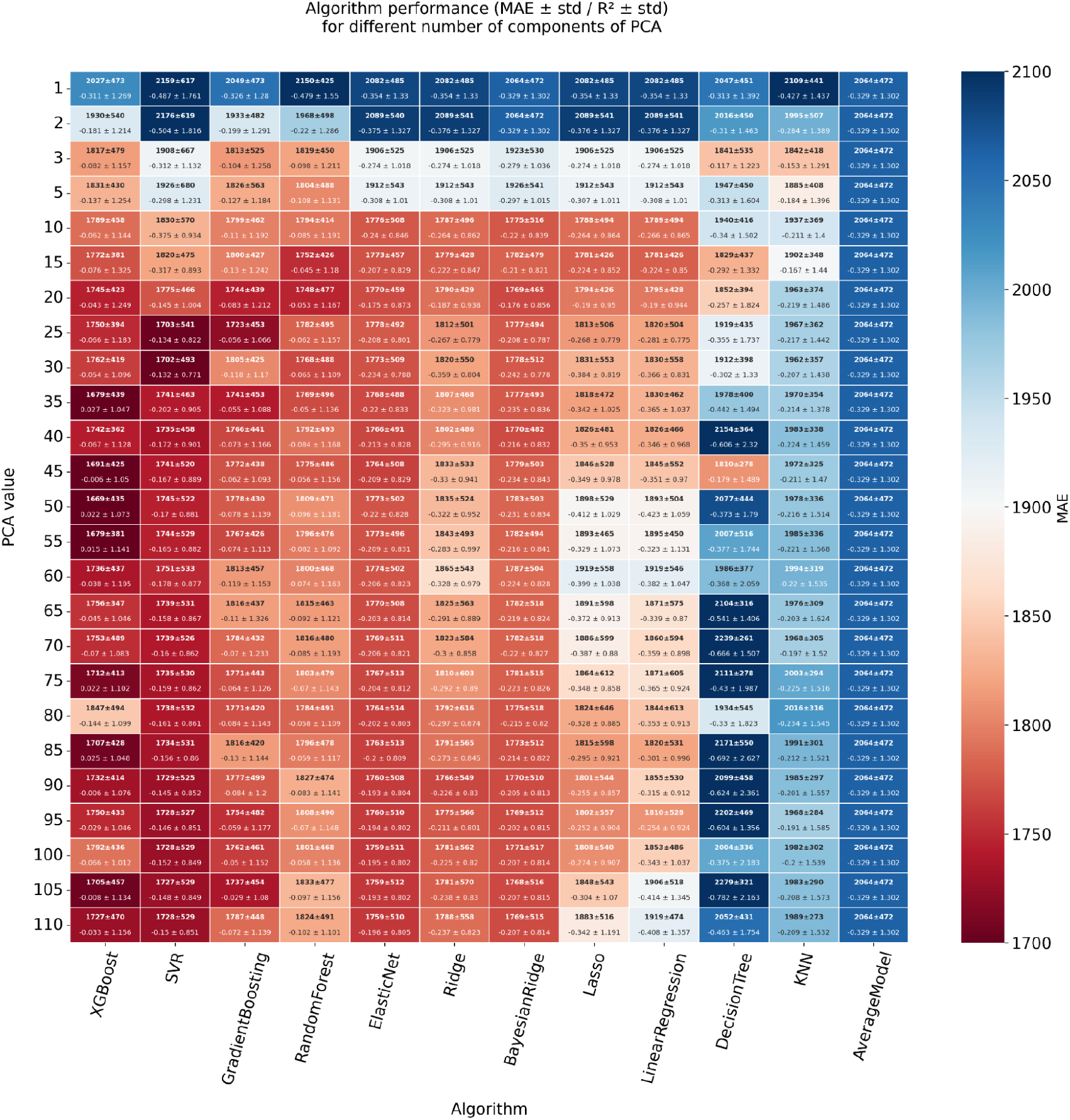
The comparison of different supervised models for predicting the age of the sample.

## Results

On Figure 3, the results of the algorithm comparison are presented. The best-performing model was XGBoost, achieving a mean absolute error (MAE) of 1669*±*435 years under 5-fold cross-validation, using 50 principal components as input features. The benchmark regressor, which predicts the mean age of the training set, achieved an MAE of 2064 *±* 472 years. Although certain models and PCA configurations produced slightly lower MAEs, these differences were generally within one standard deviation. We also evaluated the selected model on the held-out validation subset, obtaining similar results. Together, these observations indicate that the apparent performance differences are not statistically meaningful and are likely driven by noise rather than a true predictive signal. Thus, we conclude that no strong age-predictive signal is present in the training data.

The final performance on the internal Harvard test set and the external dataset is summarized in Table 1. Consistent with the cross-validation and validation results, the model fails to predict sample ages in both test settings, confirming that the age signal cannot be reliably recovered from the available features.

**Table 1:**
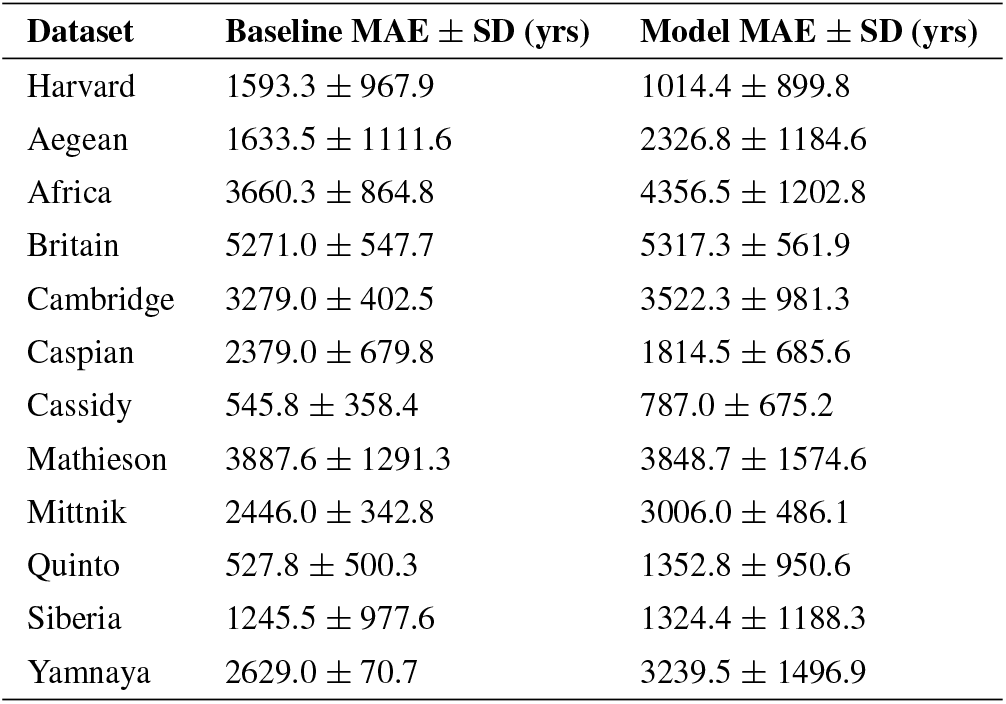
Absolute age prediction error (years) for baseline mean predictor vs. model across datasets.

**Table 2:**
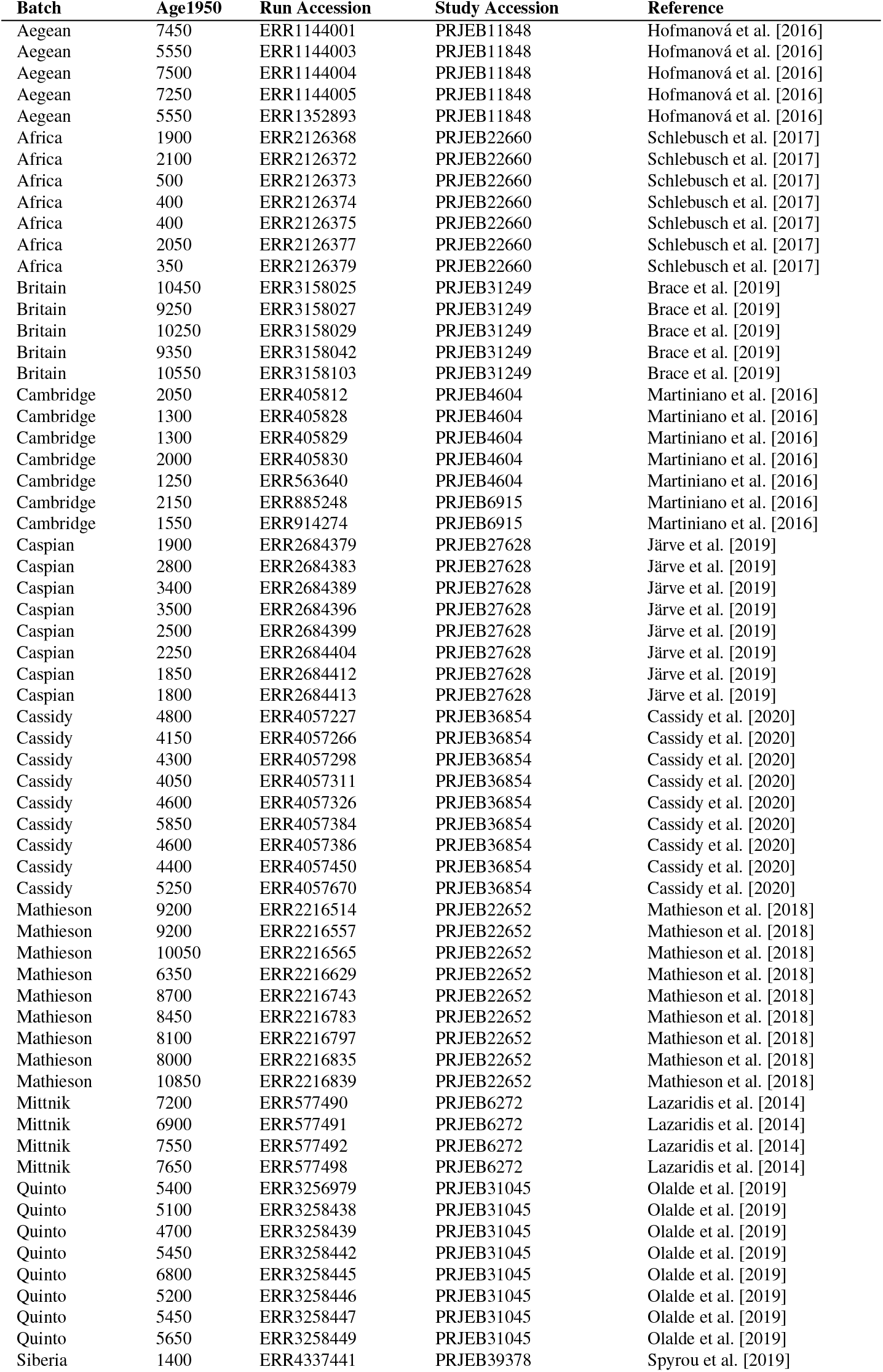
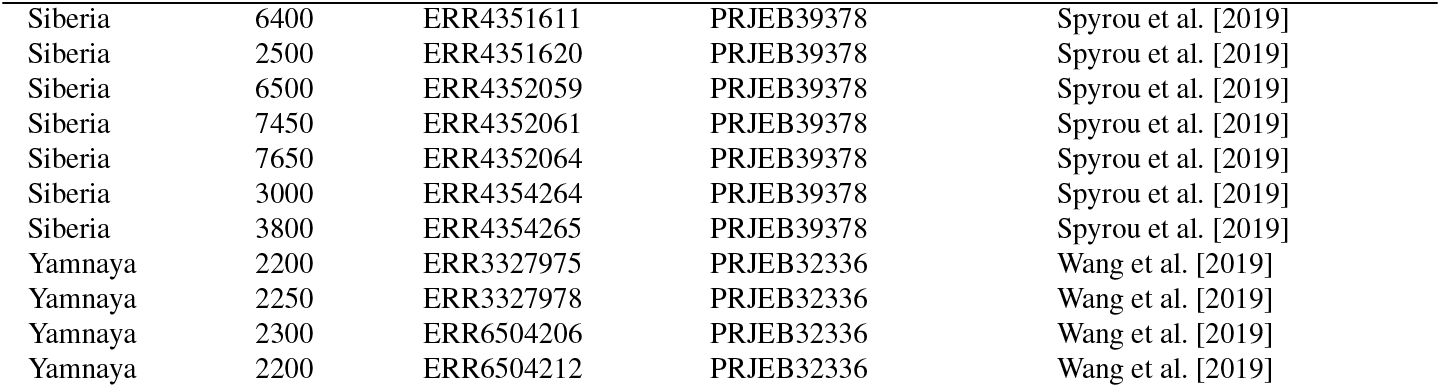
Ancient DNA samples used in this study.

| Batch | Age1950 | Run Accession | Study Accession | Reference |
| --- | --- | --- | --- | --- |
| Aegean | 7450 | ERR1144001 | PRJEB11848 | Hofmanová et al. [2016] |
| Aegean | 5550 | ERR1144003 | PRJEB11848 | Hofmanová et al. [2016] |
| Aegean | 7500 | ERR1144004 | PRJEB11848 | Hofmanová et al. [2016] |
| Aegean | 7250 | ERR1144005 | PRJEB11848 | Hofmanová et al. [2016] |
| Aegean | 5550 | ERR1352893 | PRJEB11848 | Hofmanová et al. [2016] |
| Africa | 1900 | ERR2126368 | PRJEB22660 | Schlebusch et al. [2017] |
| Africa | 2100 | ERR2126372 | PRJEB22660 | Schlebusch et al. [2017] |
| Africa | 500 | ERR2126373 | PRJEB22660 | Schlebusch et al. [2017] |
| Africa | 400 | ERR2126374 | PRJEB22660 | Schlebusch et al. [2017] |
| Africa | 400 | ERR2126375 | PRJEB22660 | Schlebusch et al. [2017] |
| Africa | 2050 | ERR2126377 | PRJEB22660 | Schlebusch et al. [2017] |
| Africa | 350 | ERR2126379 | PRJEB22660 | Schlebusch et al. [2017] |
| Britain | 10450 | ERR3158025 | PRJEB31249 | Brace et al. [2019] |
| Britain | 9250 | ERR3158027 | PRJEB31249 | Brace et al. [2019] |
| Britain | 10250 | ERR3158029 | PRJEB31249 | Brace et al. [2019] |
| Britain | 9350 | ERR3158042 | PRJEB31249 | Brace et al. [2019] |
| Britain | 10550 | ERR3158103 | PRJEB31249 | Brace et al. [2019] |
| Cambridge | 2050 | ERR405812 | PRJEB4604 | Martiniano et al. [2016] |
| Cambridge | 1300 | ERR405828 | PRJEB4604 | Martiniano et al. [2016] |
| Cambridge | 1300 | ERR405829 | PRJEB4604 | Martiniano et al. [2016] |
| Cambridge | 2000 | ERR405830 | PRJEB4604 | Martiniano et al. [2016] |
| Cambridge | 1250 | ERR563640 | PRJEB4604 | Martiniano et al. [2016] |
| Cambridge | 2150 | ERR885248 | PRJEB6915 | Martiniano et al. [2016] |
| Cambridge | 1550 | ERR914274 | PRJEB6915 | Martiniano et al. [2016] |
| Caspian | 1900 | ERR2684379 | PRJEB27628 | Järve et al. [2019] |
| Caspian | 2800 | ERR2684383 | PRJEB27628 | Järve et al. [2019] |
| Caspian | 3400 | ERR2684389 | PRJEB27628 | Järve et al. [2019] |
| Caspian | 3500 | ERR2684396 | PRJEB27628 | Järve et al. [2019] |
| Caspian | 2500 | ERR2684399 | PRJEB27628 | Järve et al. [2019] |
| Caspian | 2250 | ERR2684404 | PRJEB27628 | Järve et al. [2019] |
| Caspian | 1850 | ERR2684412 | PRJEB27628 | Järve et al. [2019] |
| Caspian | 1800 | ERR2684413 | PRJEB27628 | Järve et al. [2019] |
| Cassidy | 4800 | ERR4057227 | PRJEB36854 | Cassidy et al. [2020] |
| Cassidy | 4150 | ERR4057266 | PRJEB36854 | Cassidy et al. [2020] |
| Cassidy | 4300 | ERR4057298 | PRJEB36854 | Cassidy et al. [2020] |
| Cassidy | 4050 | ERR4057311 | PRJEB36854 | Cassidy et al. [2020] |
| Cassidy | 4600 | ERR4057326 | PRJEB36854 | Cassidy et al. [2020] |
| Cassidy | 5850 | ERR4057384 | PRJEB36854 | Cassidy et al. [2020] |
| Cassidy | 4600 | ERR4057386 | PRJEB36854 | Cassidy et al. [2020] |
| Cassidy | 4400 | ERR4057450 | PRJEB36854 | Cassidy et al. [2020] |
| Cassidy | 5250 | ERR4057670 | PRJEB36854 | Cassidy et al. [2020] |
| Mathieson | 9200 | ERR2216514 | PRJEB22652 | Mathieson et al. [2018] |
| Mathieson | 9200 | ERR2216557 | PRJEB22652 | Mathieson et al. [2018] |
| Mathieson | 10050 | ERR2216565 | PRJEB22652 | Mathieson et al. [2018] |
| Mathieson | 6350 | ERR2216629 | PRJEB22652 | Mathieson et al. [2018] |
| Mathieson | 8700 | ERR2216743 | PRJEB22652 | Mathieson et al. [2018] |
| Mathieson | 8450 | ERR2216783 | PRJEB22652 | Mathieson et al. [2018] |
| Mathieson | 8100 | ERR2216797 | PRJEB22652 | Mathieson et al. [2018] |
| Mathieson | 8000 | ERR2216835 | PRJEB22652 | Mathieson et al. [2018] |
| Mathieson | 10850 | ERR2216839 | PRJEB22652 | Mathieson et al. [2018] |
| Mittnik | 7200 | ERR577490 | PRJEB6272 | Lazaridis et al. [2014] |
| Mittnik | 6900 | ERR577491 | PRJEB6272 | Lazaridis et al. [2014] |
| Mittnik | 7550 | ERR577492 | PRJEB6272 | Lazaridis et al. [2014] |
| Mittnik | 7650 | ERR577498 | PRJEB6272 | Lazaridis et al. [2014] |
| Quinto | 5400 | ERR3256979 | PRJEB31045 | Olalde et al. [2019] |
| Quinto | 5100 | ERR3258438 | PRJEB31045 | Olalde et al. [2019] |
| Quinto | 4700 | ERR3258439 | PRJEB31045 | Olalde et al. [2019] |
| Quinto | 5450 | ERR3258442 | PRJEB31045 | Olalde et al. [2019] |
| Quinto | 6800 | ERR3258445 | PRJEB31045 | Olalde et al. [2019] |
| Quinto | 5200 | ERR3258446 | PRJEB31045 | Olalde et al. [2019] |
| Quinto | 5450 | ERR3258447 | PRJEB31045 | Olalde et al. [2019] |
| Quinto | 5650 | ERR3258449 | PRJEB31045 | Olalde et al. [2019] |
| Siberia | 1400 | ERR4337441 | PRJEB39378 | Spyrou et al. [2019] |
| Siberia | 6400 | ERR4351611 | PRJEB39378 | Spyrou et al. [2019] |
| Siberia | 2500 | ERR4351620 | PRJEB39378 | Spyrou et al. [2019] |
| Siberia | 6500 | ERR4352059 | PRJEB39378 | Spyrou et al. [2019] |
| Siberia | 7450 | ERR4352061 | PRJEB39378 | Spyrou et al. [2019] |
| Siberia | 7650 | ERR4352064 | PRJEB39378 | Spyrou et al. [2019] |
| Siberia | 3000 | ERR4354264 | PRJEB39378 | Spyrou et al. [2019] |
| Siberia | 3800 | ERR4354265 | PRJEB39378 | Spyrou et al. [2019] |
| Yamnaya | 2200 | ERR3327975 | PRJEB32336 | Wang et al. [2019] |
| Yamnaya | 2250 | ERR3327978 | PRJEB32336 | Wang et al. [2019] |
| Yamnaya | 2300 | ERR6504206 | PRJEB32336 | Wang et al. [2019] |
| Yamnaya | 2200 | ERR6504212 | PRJEB32336 | Wang et al. [2019] |

For the test dataset we used a test subset of the Harvard Dataset and additional samples outside of the Harvard dataset. We shall note that we did not include the test dataset in the cross-validation, so the test dataset was unseen for the algorithms. The results for each datapoint are presented in Appendix 1. For the Harvard dataset the results are the following. We calculated the average actual age as the average age for the test dataset. We calculated the MAE (Predicted vs Actual) as the difference between the predicted and actual age; the baseline MAE as the difference between the predicted and the average actual age.

We can see that the MAE (predicted) is more than baseline benchmark which indicates that the model does not do well on test data (which is part of the Harvard).

## Discussion

The supervised learning methods applied in this study did not produce a statistically significant improvement over the benchmark model. Although XGBoost achieved the lowest cross-validation MAE (1669 ± 435 years) with 50 PCA components, this value lies within one standard deviation of the benchmark regressor (2064 ± 472 years). The overlapping performance across algorithms and PCA configurations indicates that the observed differences are likely caused by random variation rather than a genuine signal.

These findings were further supported by the results on the independent test set, which included both Harvard dataset test samples and external data. Model predictions showed no consistent improvement over baseline estimates derived from the mean age, suggesting limited generalization beyond the training distribution.

Interestingly, when batch effects were not addressed specifically (when data from different publications were mixed during training and evaluation),the models exhibited substantially lower prediction error and appeared to capture a strong signal. Subsequent analysis, however, demonstrated that this improvement was primarily driven by batch-specific artifacts rather than by meaningful biological patterns. In other words, the models effectively learned to distinguish the source article rather than prediction the true chronological age of the samples. This observation underscores the presence of strong confounding factors tied to data origin and highlights the importance of rigorous batch control and cross-validation across studies.

However, even after applying batch-correction techniques, the confounding signal persisted, and performance did not improve, suggesting that current batch-adjustment methods are insufficient for this task and that deeper biases may be inherent in the dataset. Overall, the present approach which is based solely on statistical features and PCA-based dimensionality reduction appears insufficient for capturing a robust age-related signal in the current dataset once batch effects are properly controlled. Nonetheless, this outcome does not preclude the possibility of success with alternative strategies. Future work may benefit from incorporating domain-specific biological features, exploring non-statistical representations, or integrating complementary data sources. Additionally, expanding the dataset size could help distinguish the true signal from noise and improve model robustness.

In addition to genetic features, we explored the use of meteorological data as auxiliary input. Previous work Allentoft et al. [2012] has demonstrated that environmental conditions can influence DNA degradation rates. To account for this, we incorporated average precipitation and temperature values into our pipeline. However, we observed a strong clustering effect in these variables across samples (Figure 5). As a consequence, meteorological features became highly predictive when samples were randomly shuffled rather than grouped by geographic or climatic origin, indicating that the signal was driven by data stratification rather than biological relevance. Therefore, incorporating meteorological data into our model was determined to be infeasible for this study.

Our method produced largely negative results across multiple batches. From a biophysical perspective, one likely explanation for the inability to predict the age of the samples lies in the dynamic nature of climate: DNA degradation is driven not by a steady process, but by daily and seasonal temperature fluctuations, humidity variations, and long-term environmental shifts. These dynamic conditions mask a clear relationship between molecular damage and chronological time. Nevertheless, this does not imply that the task is impossible. With larger, cleaner datasets and improved control over environmental and technical variability (proper batch-effect), it may become feasible to extract reliable temporal signals from DNA damage profiles in future research.

## Conclusion

Our study evaluated the potential of using machine learning to estimate the chronological age of ancient DNA samples based on their molecular damage profiles. Once batch effects were controlled, the apparent predictive signal largely disappeared, indicating that experimental and environmental factors dominate the variation in DNA damage patterns. None of the tested algorithms consistently outperformed a simple mean-based baseline.

These findings suggest that damage profiles alone are insufficient for reliable age estimation. However, they also highlight important directions for future research. Incorporating additional biological and contextual information such as preservation environment, biochemical parameters, and physical models of DNA decay could improve predictive power. Expanding dataset size and diversity may further help distinguish genuine chronological signals from background noise. Thus, while our current approach did not achieve accurate molecular dating, it provides an benchmark and methodological insight for subsequent developments in the field.

All code used in this work is publicly available at: https://github.com/maksimkazanskii/DNA_age_prediction.

## Acknowledgements

We thank O. Limanovskaya, O. Mozhey, and O. Kantidze for valuable discussions and support. We used large language model (LLM) assistance (OpenAI GPT-5) to help polish the language and improve clarity of the manuscript. All data analysis, scientific interpretations, and conclusions were generated by the authors.

## Appendix A Ancient DNA Samples Used in This Study

### Column Descriptions

**Batch** Study or dataset identifier indicating the publication or source group to which the sample belongs.

**Age1950** Estimated age of the individual in years before 1950 (the standard archaeological reference year).

**Run Accession** Sequence run identifier from public archives (e.g., ENA/SRA) corresponding to the specific BAM file.

**Study Accession** Project-level accession identifier for the study in which the sample was generated.

**Reference** Citation for the original publication reporting and describing the ancient DNA data.

## Notes

### Competing Interest Statement

The authors have declared no competing interest.

